# Structural insights into Charcot-Marie-Tooth disease-linked mutations in human GDAP1

**DOI:** 10.1101/2022.02.18.481076

**Authors:** Aleksi Sutinen, Giang Thi Tuyet Nguyen, Arne Raasakka, Gopinath Muruganandam, Remy Loris, Emil Ylikallio, Henna Tyynismaa, Luca Bartesaghi, Salla Ruskamo, Petri Kursula

**Affiliations:** Faculty of Biochemistry and Molecular Medicine & Biocenter Oulu, University of Oulu, Finland; Department of Biomedicine, University of Bergen, Norway; VIB-VUB Center for Structural Biology, Vlaams Instituut voor Biotechnologie, Brussels, Belgium; Structural Biology Brussels, Department of Bioengineering Sciences, Vrije Universiteit Brussel, Belgium; Stem Cells and Metabolism Research Program, Faculty of Medicine, University of Helsinki, Finland; Clinical Neurosciences, Neurology, Helsinki University Hospital, Finland; Department of Neuroscience, Karolinska Institutet, Sweden

**Keywords:** Charcot-Marie-Tooth disease, GDAP1, GST superfamily, protein structure, neuropathy

## Abstract

Charcot-Marie-Tooth disease (CMT) is the most common inherited peripheral polyneuropathy in humans, and its different subtypes are linked to mutations in dozens of different genes. Mutations in ganglioside-induced differentiation-associated protein 1 (GDAP1) cause two types of CMT, demyelinating CMT4A and axonal CMT2K. The GDAP1-linked CMT genotypes are mainly missense point mutations. Despite clinical profiling and *in vivo* studies on the mutations, the etiology of GDAP1-linked CMT is poorly understood. Here, we describe the biochemical and structural properties of the Finnish founding CMT2K mutation H123R as well as CMT2K-linked R120W, both of which are autosomal dominant mutations. The disease variant proteins retain close to normal structure and solution behaviour, but both present a large decrease in thermal stability. Using GDAP1 variant crystal structures, we identify a side chain interaction network between helices α3, α6, and α7, which is affected by CMT mutations, as well as a hinge in the long helix α6, which is linked to structural flexibility. Structural analysis of GDAP1 indicates that CMT may arise from disruption of specific intra- and intermolecular interaction networks, leading to alterations in GDAP1 structure and stability, and eventually, insufficient motor and sensory neuron function.

---

Inherited polyneuropathies are a genetically and clinically diverse group of neurodegenerative diseases, which affect motor and sensory neurons in the peripheral nervous system (PNS) [1, 2]. Mutations in dozens of genes expressed in the PNS cause Charcot-Marie-Tooth syndrome (CMT). Based on clinical findings, CMT can be classified into three forms: demyelinating, axonal, and intermediate [3, 4]. The progress of CMT is linked to the hereditary pattern, whereby the autosomal recessive form has an earlier onset and more severe symptoms than the autosomal dominant form [5–7]. Understanding the molecular function of the proteins involved in the etiology of neuropathies is vital in efforts towards treatment and diagnosis.

Ganglioside-induced differentiation-associated protein 1 (GDAP1) is an integral mitochondrial outer membrane (MOM) protein, and the *GDAP1* gene is one of the most abundant in missense mutations linked to CMT [8–10]. Both autosomal dominant and recessive modes of inheritance are found, resulting in either autosomal recessive, or dominant demyelinating CMT4, autosomal dominant axonal CMT2 or intermediate CMTRIA types of CMT, with varying phenotype severity [11]. The mutations R120W and H123R, which we focus on in this study, are both autosomal dominant mutations causing the CMT2K subtype. Both phenotypes show typical slow development after onset, and main symptoms include loss of sensation in limb extremities and muscle weakness. The clinical profiling of the phenotypes has been described earlier in Spain and Finland [12–14]. GDAP1 is ubiquitously expressed in tissues, but most of the expression confines to neuronal tissues [9, 15]. In the cell, GDAP1 localizes as a tail-anchored MOM protein [16]. Structurally, GDAP1 resembles glutathione *S*-transferases (GST), and it contains unique flexible loops [17, 18]. The most accurate structural data thus far cover the dimeric core GST-like domain of human GDAP1, including the GDAP1-specific insertion [18]. The transmembrane helix and the GST-like domain are linked by a hydrophobic domain and possibly a flexible linker loop.

GST superfamily members function in prokaryotic and eukaryotic metabolism through the utilization of reduced glutathione to catalyse a range of chemically diverse reactions. GSTs often contribute to mechanisms of neurodegenerative diseases [19, 20]. In comparison to many other enzyme superfamilies, GSTs are unique in that sequence conservation appears to be driven by fold stability instead of catalytic features, as reflected in the broad spectrum of GST substrates [21, 22]. While the function of GDAP1 is not fully understood at the molecular level, it has been linked to multiple mitochondrial events in neurons [23, 24], redox regulation, and signal transduction [25, 26].

In the Finnish population, the autosomal dominant founder mutation H123R accounts for as much as 20-30% of the local CMT cases [12, 27]. We carried out structural analysis of two selected autosomal dominant GDAP1 mutants, H123R and R120W, using X-ray crystallography and complementary biophysical and computational techniques. In addition, we used three cell culture models, rat dorsal root ganglion (rDRG) neurons, human embryonic kidney 293 (HEK-293T) cells, and human skin fibroblasts, to observe the oligomeric state of GDAP1 and the effects of the disease mutations therein.

## MATERIALS AND METHODS

### Cloning

The GDAP1Δ295-358 and GDAP1Δ303-358 constructs used to produce soluble recombinant human GDAP1 in *E. coli* have been described [18]. Point mutations were generated in GDAP1Δ303-358 by a site-directed mutagenesis protocol with *Pfu* polymerase [28]. An N-terminal His_6_-affinity tag and a Tobacco Etch Virus (TEV) protease digestion site were included in each construct. The full-length GDAP1 coding sequence was subcloned into Gateway^®^ (Invitrogen) vectors pEN-TTmcs and pSLIK-HYGRO [29]. Point mutations were introduced as above. In addition to the *GDAP1* gene, the tetracycline-responsive promoter element (tight-TRE) was added within the cloning site [30], and a single N-terminal FLAG-tag was introduced into each construct [31]. All constructs were verified by DNA sequencing.

### Recombinant protein production

Soluble recombinant GDAPΔ295-358 and GDAP1Δ303-358 were expressed in *E. coli* BL21(DE3) strain using ZYM-5052 autoinduction medium (24 h, 220 rpm, +37 °C) [32]. The cells were re-suspended in binding buffer (40 mM HEPES, 400 mM NaCl, 2% glycerol, and 25 mM imidazole (pH 7.5)), containing EDTA-free protease inhibitor tablet (Sigma), snap-frozen in liquid nitrogen and stored at −70 °C. Lysis of the cells was done by sonication, and the lysate was clarified by centrifugation (40 min, 16 000 rpm, +4 °C). Recombinant protein was captured on a Ni^2+^-NTA HisPur^®^ affinity resin by gravity flow (Thermo Fisher Scientific). Unbound proteins were washed with binding buffer. The matrix was eluted with the same buffer, with imidazole at 250 mM. The affinity tag was cleaved with TEV protease treatment in 25 mM HEPES, 300 mM NaCl, 2% glycerol, 1 mM TCEP in a dialysis tube (16 h, +4 °C). The His_6_ -tag and TEV protease were then removed by another Ni^2+^-NTA affinity step. Size exclusion chromatography (SEC) was performed on a Superdex 75 10/300 GL increase column (Cytiva) using 25 mM HEPES (pH 7.5), 300 mM NaCl (SEC buffer) as mobile phase.

For GDAPΔ295-358, the Ni^2+^-NTA purification protocol was identical, but 40 mM HEPES, 400 mM NaCl, 20 mM imidazole, pH 7.5 was used as lysis and Ni-NTA washing buffer, and 32 mM HEPES, 320 mM NaCl, 500 mM imidazole, pH 7.5 was used to elute bound proteins. EDTA-free protease inhibitor cocktail (Roche) was included during cell freezing and lysis. TEV protease treatment was performed in dialysis against 40 mM HEPES, 400 mM NaCl, pH 7.5 at +4 °C overnight, followed by a second Ni^2+^-NTA affinity step. SEC was performed using a Superdex 200 16/60 HiLoad column (Cytiva) with 20 mM HEPES, 300 mM NaCl, 1% (v/v) glycerol, 0.5 mM TCEP, pH 7.5 as mobile phase.

SEC peak fractions were analyzed with SDS-PAGE, and Coomassie-stained bands were used for protein identification using a Bruker UltrafleXtreme matrix-assisted laser desorption/time-of-flight mass spectrometer (MALDI TOF-MS). Tryptic peptides extracted from the gel were identified by a search in NBCI and SwissProt databases using BioTools2.2 (Bruker).

### Crystallization, data collection, and structure determination

Mutant GDAP1Δ303-358 crystals were obtained using the sitting-drop vapor diffusion method at +4 °C. Proteins were mixed with mother liquor on crystallization plates using a Mosquito LCP nano-dispenser (TTP Labtech). The protein concentration was 10-30 mg/ml in 75 nl, and 150 nl of reservoir solution were added. H123R crystals were obtained in 0.15 M *DL*-malic acid, 20% (w/v) PEG3350. R120W crystals were obtained in 0.1 M HEPES (pH 7.3) and 10% (w/v) PEG6000. Crystals were briefly soaked in a mixture containing 10% PEG200, 10% PEG400, and 30% glycerol for cryo-protection, before flash freezing in liquid N_2_.

A novel crystal form of wild-type GDAP1Δ295-358 was obtained at +8 °C in 200 mM NH4 formate, 25% PEG3350 in a drop containing 150 nl of 8.64 mg/ml protein and 150 nl of reservoir solution. Cryoprotection was performed by adding 3 μl of cryoprotectant solution (75% (v/v) reservoir solution mixed with 25% (v/v) PEG200) directly into the crystallization drop, followed by crystal mounting and flash freezing with liquid N_2_.

Diffraction data collection at 100 K was conducted at the PETRA III synchrotron source (DESY, Hamburg, Germany), on the P11 beamline [33, 34] and the EMBL/DESY P13 beamline. Diffraction data were processed and scaled using XDS [35]. Crystal structures of wild-type GDAP1Δ303-358 [18] were used as search models in molecular replacement (MR). MR, model refinement, and structure validation were done using Phenix [36, 37] and CCP4 [38]. The models were refined using Phenix.Refine [39] and rebuilt using COOT [40]. The structures were validated using MolProbity [41]. The data processing and structure refinement statistics are in **Table I**, and the refined coordinates and structure factors were deposited at the Protein Data Bank with entry codes 7Q6K (R120W), 7Q6J (H123R), and 7YWD (new crystal form of wtGDAP1). The diffraction datasets for the mutants were uploaded on Zenodo: https://doi.org/10.5281/zenodo.4686880 (R120W) and https://doi.org/10.5281/zenodo.4686876 (H123R).

**Table 1.** Crystallographic data processing and structure refinement. Data for the highest-resolution shell are shown in parentheses.

| Protein | R120W GDAP1 | H123R GDAP1 | wtGDAP1 |
| --- | --- | --- | --- |
| *Data collection* |  |  |  |
| Beamline | P11/PETRA III | P11/PETRA III | P13 |
| X-ray wavelength (Å) | 1.0332 | 1.0332 | 1.0332 |
| Space group | P2<sub>1</sub>2<sub>1</sub>2<sub>1</sub> | P6<sub>3</sub>22 | P3<sub>1</sub>21 |
| Unit cell dimensions a, b, c (Å) | 72.71 115.88 116.18 | 147.27 147.27 114.56 | 126.8 126.8 177.1 |
| Resolution range (Å) | 50 - 2.2 (2.3 - 2.2) | 50 - 3.4 (3.5 - 3.4) | 100 - 3.2 (3.39 – 3.20) |
| Completeness (%) | 99.7 (98.8) | 99.5 (98.6) | 99.9 (99.9) |
| Redundancy | 6.5 (6.7) | 12.8 (12.6) | 9.8 (9.9) |
| R<sub>meas</sub> (%) | 9.0 (191.3) | 41.1 (339.8) | 10.7 (387.4) |
| <I/σI> | 12.8 (1.0) | 6.7 (0.9) | 16.0 (0.6) |
| CC<sub>1/2</sub> (%) | 99.9 (69.6) | 99.6 (60.5) | 100.0 (17.7) |
| Wilson B (Å<sup>2</sup>) | 49.4 | 104.5 | 129.4 |
| *Structure refinement* |  |  |  |
| R<sub>cryst</sub>/R<sub>free</sub> (%) | 21.1/ 23.3 | 25.0 / 29.1 | 25.1 / 27.6 |
| RMSD bond lengths (Å) | 0.013 | 0.002 | 0.003 |
| RMSD bond angles (°) | 1.35 | 0.43 | 0.61 |
| Molprobity score | 1.17 | 0.91 | 2.01 |
| Ramachandran favoured/<br>outliers (%) | 95.89 / 0.6 | 95.90 / 0.82 | 95.4 / 1.6 |
| PDB entry | 7Q6K | 7Q6J | 7YWD |

### Modelling, simulation, and bioinformatics

A model for full-length GDAP1 was obtained from AlphaFold2 [42] and used for further analyses as such. In addition, crystal structure-based models were prepared and analysed. Missing loops of the wtGDAP1 crystal structure were built with YASARA [43], and the structure was minimized. The model was further used as a starting point for SAXS data fitting (see below) as well as molecular dynamics simulations.

MD simulations were run on a GDAP1 monomer, with all loops in place. The simulations were run using GROMACS [44], with input file preparation on CHARMM-GUI [45]. The force field used was CHARMM36m [46], with a cubic box with a 10 Å extension around the protein. Solvation was done with the TIP3P water model in 0.15 M NaCl. Temperature (NVT) equilibration to 300 K and pressure (NPT) equilibration, via isotropic pressure coupling, were carried out using the Berendsen thermostat.

Structural properties of GDAP1 were analysed with bioinformatics tools, including NAPS [47] for centrality analyses and DynaMine [48] for prediction of flexibility. Stability effects of missense mutations were predicted with CUPSAT [49]. Hydrophobic clusters were identified with ProteinTools [50].

### Small-angle X-ray scattering

The structure and oligomeric state of the GDAP1 R120W and H123R mutants were analysed with SEC-coupled small-angle X-ray scattering (SAXS). SEC-SAXS experiments were performed on the SWING beamline [51] (SOLEIL synchrotron, Saint Aubin, France). Samples were dialyzed against fresh SEC buffer and centrifuged at >20000 g for 10 min at +4 °C to remove aggregates. 50 μl of each protein sample at 8.5-10 mg/ml was injected onto a BioSEC3-300 column (Agilent) at a 0.2 ml/min flow rate. SAXS data were collected at +15 °C, over a q-range of 0.003–0.5 Å^-1^ (q = *4π sin(θ)/λ*, where 2θ is the scattering angle).

Further processing and modelling were done using ATSAS 3.0 [52]. Scattering curves were analysed and particle dimensions determined using PRIMUS [53] and GNOM [54], respectively. Chain-like *ab initio* models were generated using GASBOR [55]. In a complementary approach, different GDAP1 dimer models were fitted against the experimental SAXS data using CRYSOL [56]. SUPCOMB was used to superimpose SAXS models and crystal structures [57].

### Circular dichroism spectroscopy

Synchrotron radiation circular dichroism (SRCD) spectra were collected from 0.5 mg/ml samples on the AU-CD beamline at the ASTRID2 synchrotron source (ISA, Aarhus, Denmark). The samples were prepared in a buffer containing 10 mM HEPES pH 7.5, 100 mM NaF. The samples were equilibrated to room temperature and applied into 0.1-mm pathlength closed circular quartz cuvettes (Suprasil, Hellma Analytics). SRCD spectra were recorded from 170 nm to 280 nm at +25 °C. Three repeat scans per measurement were recorded and averaged. The CD spectra baselines were processed and converted to molar ellipticity using CDToolX [58].

### Thermal stability

Thermal unfolding of GDAP1 variants in SEC buffer was studied by nanoDSF using a Prometheus NT.48 instrument (NanoTemper). Tryptophan fluorescence was excited at 280 nm, and emission was recorded at 330 and 350 nm. The samples were heated from +20 to +90 °C with a heating rate of 1 °C/min, and changes in the fluorescence ratio (F350/F330) were monitored to determine apparent melting points. The data were analyzed using Origin (OriginLab Corporation, Northampton, MA, USA)

### Cell culture and Western blotting

Human skin fibroblast cultures were established from skin biopsies of a healthy donor and a patient with GDAP1 H123R mutation [12, 59]. Written consent for the use of patient material was obtained, and the study was approved by the Coordinating Ethics Committee of the Helsinki and Uusimaa Hospital District. The purification of rDRG sensory neurons, the generation of lentiviral particles, and their use to overexpress the GDAP1 constructs were done as described [60] for MORC2. rDRGs were matured into sensory neurons, and GDAP1 expression was induced by doxycycline-initiated tight-TRE promotor expression using the Lentivirus system [30]

The protein fractions were isolated from the rDRG and fibroblast cells and membranes using 40 mM HEPES pH 7.0, 400 mM NaCl, 1% n-Dodecyl-β-D-Maltopyranoside (DDM), and the supernatant was clarified by centrifugation 45 000 rpm, +4 °C. The proteins were separated with 12% SDS-PAGE under non-reducing conditions.

HEK293T-D10 cells were used to serve as endogenous control and to test the redox sensitivity of the mammalian-derived GDAP1 samples. Proteins were isolated from total cell lysate and mitochondrial fraction. The mitochondria were isolated from the cells using linear 15-50% (w/v) sucrose gradient centrifugation. The protein was treated with similar lysis conditions as above, and SDS-PAGE was performed with and without 192 mM β-mercaptoethanol.

Proteins were transferred onto 0.22 μm nitrocellulose membranes with the semi-dry transfer protocol in TurboBlot^®^-buffer (Bio-Rad Laboratories, Inc.). The membrane was blocked with Tris-buffered saline, 20 mM Tris-HCl (pH 7.4), 100 mM NaCl, 0.1% v/v Tween-20 (TBST), 5% w/v casein (milk powder) and incubated for 2 h at +4 °C. The primary antibody, rabbit anti-GDAP1 anti-serum [16] was added at a 1:5000 dilution and incubated for 1 h at +4 °C, followed by the secondary antibody for 1 h at +4 °C (anti-rabbit IgG-HRP, Promega 65-6120). The Pierce^®^ enhanced chemiluminescence substrate (Thermo-Fischer Scientific) was added, and the blot was illuminated using ChemiDoc XRS+ (Bio-Rad). Tubulin was used as a loading control in all experiments.

### Immunofluorescence microscopy

rDRG cells were fixed with 4% paraformaldehyde in phosphate buffered saline 8 mM Na_2_HPO_4_, 2 mM KH_2_PO_4_, 137 mM NaCl, 2.7 mM KCl (pH 7.4) (PBS) at +22 °C for 10 min and washed in PBS. They were then incubated 1 h at +22 °C in a blocking solution (5% bovine serum albumin, 1% goat serum, 0.3% Triton X-100 in PBS) and with primary antibodies overnight at +4 °C (primary Abs: mouse anti-flag –Sigma F1804; rabbit anti-NF-145 – Millipore AB1987), followed by washing in PBS. The secondary antibodies were incubated for 45 min at +22 °C (secondary Abs: anti-mouse 594, Invitrogen A11005; anti-rabbit 488, Invitrogen A11034). The cells were counterstained with 1:10000 4’,6-diamidino-2-phenylindole (DAPI) in PBS for 5 min at +22 °C. The fixed samples were mounted on cover slips with Vectashield, and images were acquired with a Zeiss LSM700 confocal microscope (Carl Zeiss AG).

## RESULTS

### Structural effects of CMT mutations H123R and R120W on GDAP1

The CMT-linked missense mutations in GDAP1 are clustered within the vicinity of the hydrophobic clusters of the N-terminal GST-like domain (GSTL-N), the C-terminal GST-like domain (GSTL-C), and the dimer interface. The affected side chains are often polar or charged and orient towards the solvent (**Fig. 1A**). They are also close to the hydrophobic clusters of GDAP1 (**Fig. 1B**). For example, the α6 helix, Lys188-Glu229, has 20 charged residues along the helix. The clustered mutations could change the side chain interaction networks between helices α3, α6, and α7, which further might affect GDAP1 folding and stability. Here, we focused on two CMT-linked GDAP1 mutations on helix α3 pointing towards α6, R120W and H123R (**Fig. 1A**).

**Figure 1.**
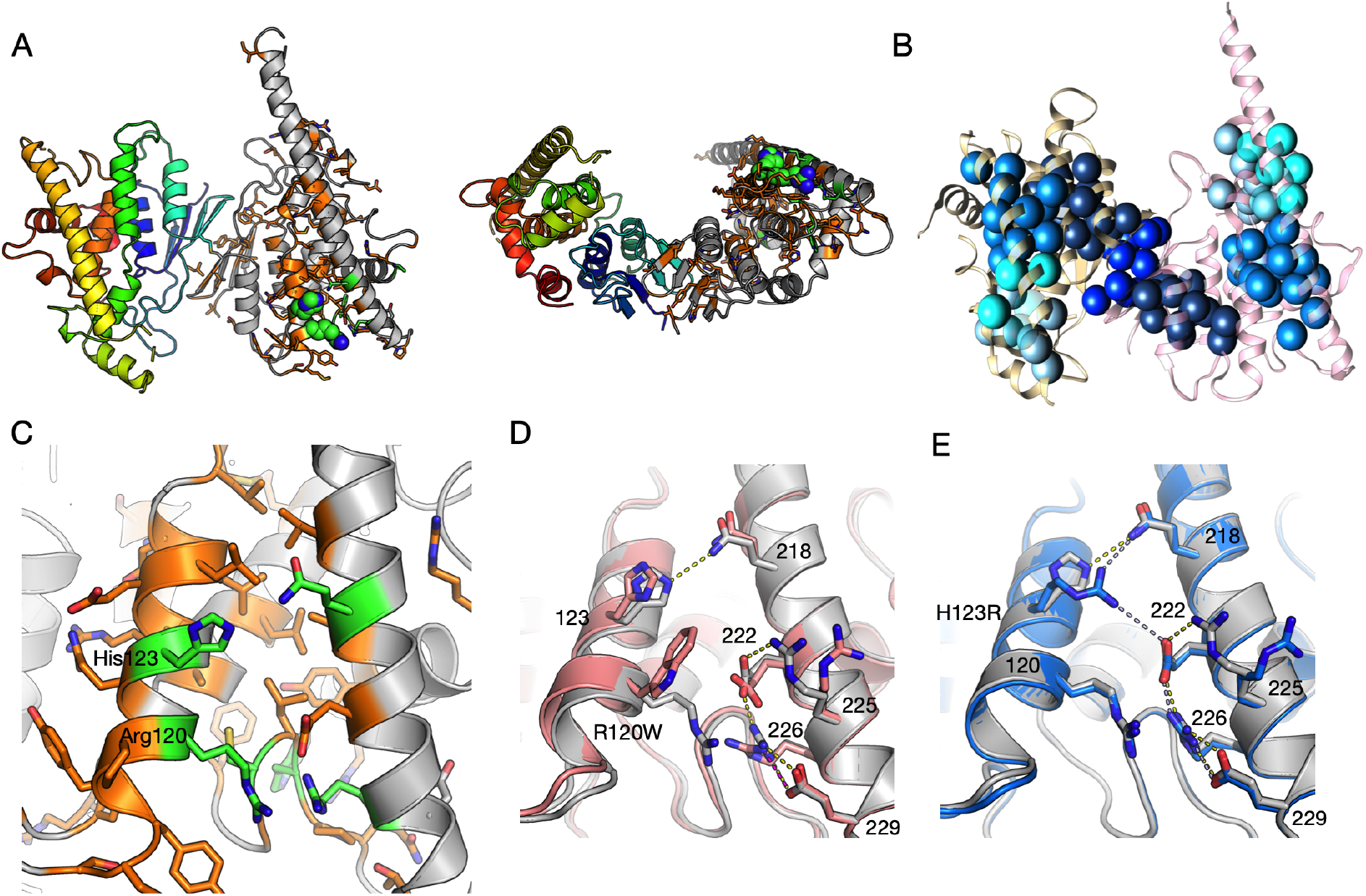
Crystal structure of GDAP1 and effects of CMT mutations. A. The overall structure of the wtGDAP1 core domain dimer in two orientations, as published before [18]. The left monomer is coloured with rainbow colours, while one on the right is gray and shows the positions of CMT mutations, with side chains visible. The mutations linked to CMT2K are green, while the rest are orange. Arg120 and His123 are shown as spheres. In the right-hand orientation, the mitochondrial outer membrane would be below the dimer. B. Hydrophobic clusters in wtGDAP1. The orientation is the same as the left panel of A, indicating that many CMT mutations lie in close vicinity of the hydrophobic cores. C. Zoom in on helices α3, α6, and α7. Colouring of CMT mutations is as in A; side chains are shown only for CMT-linked positions. Note how the CMT-linked residues participate in a large intramolecular network of interactions. D. Comparison of wtGDAP1 (gray) and R120W (pink) crystal structures. E. Comparison of wtGDAP1 (gray) and H123R (blue). Hydrogen bonds are shown as dashed lines.

We previously determined the crystal structure of wild-type GDAP1Δ303-358, which corresponds to the GST-like core domain of GDAP1 in dimeric form, including the GDAP1-specific insertion [18]. Here, we expressed and purified the variants R120W and H123R, compared them to wild-type GDAP1 (wtGDAP1) crystal and solution structures, and studied their folding and thermal stability. The crystal of the R120W mutant variant had a new crystal form, while H123R had the same space group as the wtGDAP1 structure, displaying a homodimer in the asymmetric unit. In the H123R structure, the dimer in the crystal is covalently linked *via* a disulphide bond at Cys88, like the wtGDAP1 protein [18]. The disulphide bridge *via* Cys88 also exists in the R120W structure, but the dimer is formed *via* crystallographic symmetry.

Both Arg120 and His123 are on the α3 helix, partially solvent-accessible (**Fig. 1C**). In both mutant structures, as in wtGDAP1, the most flexible regions are in loops between β3-β4 at positions Leu71-Ala77, and α5-α6 at positions Arg159-Ile186. The β3-β4 loop is more structurally ordered in the mutants compared to wtGDAP1. The α5-α6 region corresponds to the GDAP1-specific insertion in the GST superfamily [61]. In both mutant structures, the flexible loop between helices α6-α7 is similar to wtGDAP1; the Cα backbone is visible, but side chains have poor density.

In the crystal state, the mutations do not cause major structural changes (**Fig. 1D,E, S2**). However, intramolecular interactions are altered. In the H123R structure (**Fig. 1E**), the His123-Tyr124 π-orbital interaction is disturbed in the mutant, while the interaction with the side chain of Gln218 is preserved. The salt bridge network around Glu222 and Arg226 is conserved and now includes Arg123. In chain B, the electron density for Arg123 is weak, indicating flexibility of the mutant residue.

Arg120 in wtGDAP1 forms a H-bond with the backbone carbonyl of Cys240, and it is part of a salt bridge network involving Glu222, Arg226, and Glu229 (**Fig. 1C-E).** Intriguingly, Arg120 has a close contact with Arg226 in wt-GDAP1, whereby the two Arg π systems stack, and the surrounding Glu222 and Glu229 neutralize charges *via* salt bridges. Trp120, as a bulky side chain, causes steric hindrance in the R120W mutant (**Figure 1D**), and the α3 helix, carrying Trp120, moves outwards by ~1 Å, and the contact with the neighbouring α6 is weakened. The backbone interaction to Cys240 is lost in the mutant, and the salt bridge network centered at Arg226 is disturbed as is the contact between His123 and Gln218, which could be linked to loss of protein stability.

### The mutant proteins show unaltered conformation but lowered stability

To compare the solution and crystal structures, SAXS analysis was performed on the H123R and R120W mutants (**Fig. 2**). SEC-SAXS was employed to achieve better separation between monomer and dimer fractions. Previously, we showed this equilibrium to be concentration-dependent; high concentration favours the dimeric form [18]. The SEC-SAXS profiles show that the samples are monodisperse with a same radius of gyration, R_g_, across the peak **(Fig. S1)**. The molecular weight across the main peak showed that in both mutants, the peak contained a dimeric form similar to wtGDAP1; accordingly, the scattering curves were essentially identical.

**Figure 2.**
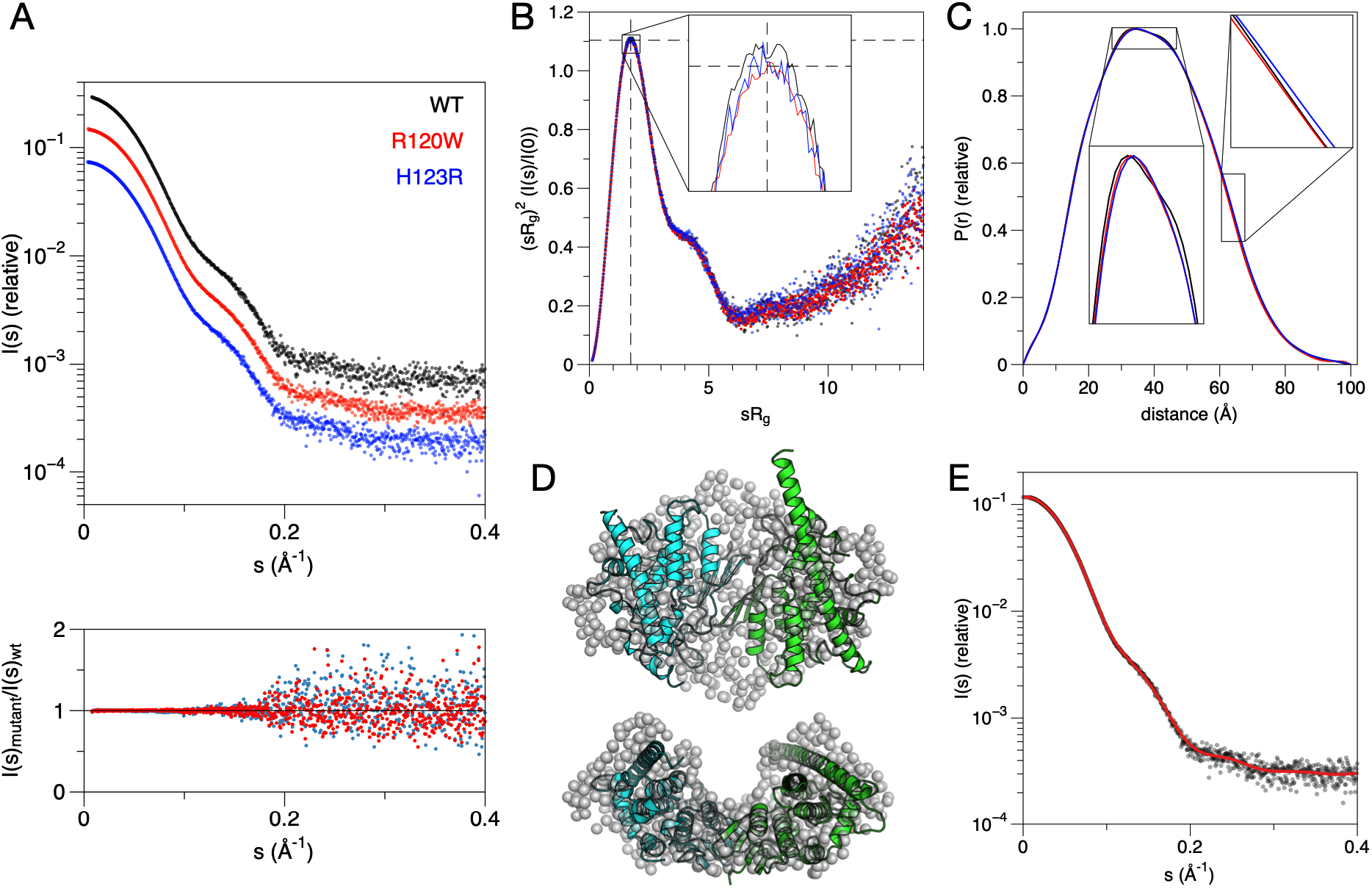
SAXS analysis of the GDAP1 mutations. A. Top: Scattering curves from synchrotron SEC-SAXS. The curves are diplaced in the y direction for clarity. Bottom: comparison of the mutant data to wtGDAP1 indicates that the SAXS data are essentially identical. B. Dimensionless Kratky plot shows clobular structure and same level of flexibility. The dashed lines crossing (x=√3, y=1.1) reflects the theoretical maximum for a rigid globular particle. C. Distance distributions indicate similar size and shape, with ver minor differences when zoomed in. D. *Ab initio* chain-like model (gray spheres) overlaid with the crystal structure of wtGDAP1 [18]. Note how the long helix α6 from the extended conformation does not fit into the envelope. E. Fit of the *ab initio* model in panel D (red line) to the SAXS data from wtGDAP1 (black dots).

Further analysis revealed that the R_g_ values matched the ones for wtGDAP1 (**Table II**), indicating similar solution conformation and oligomeric state. Both mutants possess a similar globular fold with essentially the same level of flexibility as wtGDAP1, as demonstrated by dimensionless Kratky plots (**Fig. 2B**). Distance distribution functions revealed that the particle dimensions in solution are nearly identical between the mutants and wtGDAP1 (**Fig. 2C**). Hence, at the resolution of a SAXS experiment, neither mutation caused large-scale conformational changes.

**Table 2.** SAXS parameters. The values for wtGDAP1 are from our previous study [18].

| Construct | GDAP1Δ303-358 H123R | GDAP1Δ303-358 R120W | GDAP1Δ303-358 wt |
| --- | --- | --- | --- |
| $R_g$ (Å) from P(r) | 30.74 ± 0.05 | 30.64 ± 0.05 | 30.60 ± 0.11 |
| $R_g$ (Å) from Guinier plot | 30.73 | 30.64 | 30.70 ± 0.03 |
| $D_{max}$ (Å) | 99.9 | 99.95 | 99.00 |
| Porod volume estimate, $V_p$ (Å<sup>3</sup>) | 107157.00 | 107947 | 105,750 |
| s$R_g$ limits | 0.24-1.29 | 0.24-1.30 | 0.41-1.30 |
| MW from consensus Bayesian assessment based on SAXS data (kDa) | 74.3 | 74.3 | 72.4 |
| Calculated monomeric MW from sequence (kDa) | 35.1 | 35.1 | 35.2 |
| SASBDB entry | XXX | XXX | SASDJV8 |

The dimer-monomer equilibrium in GDAP1 is dynamic, and the dimeric form is favoured at high concentrations [18]. At the concentrations and conditions used here, GDAP1 exists as a dimer, as indicated by the SAXS data. Chain-like *ab initio* models confirmed the observation that both mutants are nearly indistinguishable from dimeric wtGDAP1 (**Fig. 2D,E**). Thus, there is no indication of effects on oligomeric status by the two mutations.

The SAXS data indicate a dimeric form for both mutants (**Table II**). The ambiguity between the two mutants was estimated with AMBIMETER [62], and H123R and R120W have both similar levels of globularity and stability. These observations are in line with the fairly minor conformational differences in the crystal state, whereby R120W – as a more drastic replacement - led to a small movement of the α3 helix and loss of hydrogen bonding interactions. On the other hand, the only fully monomeric GDAP1 mutant we studied earlier, Y29E/C88A, is more globular than any of the dimeric forms [18].

To further compare the molecules in solution and in the crystal, the crystal structure coordinates were fitted against the SAXS scattering curve (**Fig. 2D**). Based on the analysis, it is obvious that both mutants adopt the dimer form, having very similar folds in solution as the wtGDAP1. After building in the loops, the dimer structure fits to the data better than the crystal structure (**Fig. S3**); hence, the conformation observed in the crystal represents the solution structure, with the addition of the GDAP1-specific flexible insertion.

Since the mutants presented similar solubility and folding as wtGDAP1, we tested whether the mutations cause changes to GDAP1 stability or secondary structure content. To test for quantitative differences in secondary structures, we measured SRCD spectra (**Fig. 3A**). The SRCD spectra of the mutants overlay well with the wtGDAP1 CD spectrum, showing that the mutants on average have a similar secondary structure composition in solution as wtGDAP1. The CD peak at 208 nm is weaker for H123R, which may indicate minor differences in intramolecular interactions between α-helices, as seen in the crystal structure. Once the GDAP1 dimer forms *via* the disulfide bond, the structure becomes very stable, and it is challenging to dissociate the dimer [18]. Due to the high helical content in GDAP1 and the similarity of the CD spectra, we can confirm that the mutations do not affect the overall folding characteristics of GDAP1. Small differences in spectral shape may be caused by both local stacking of amino acid side chains as well as interactions between secondary structure elements [63, 64].

**Figure 3.**
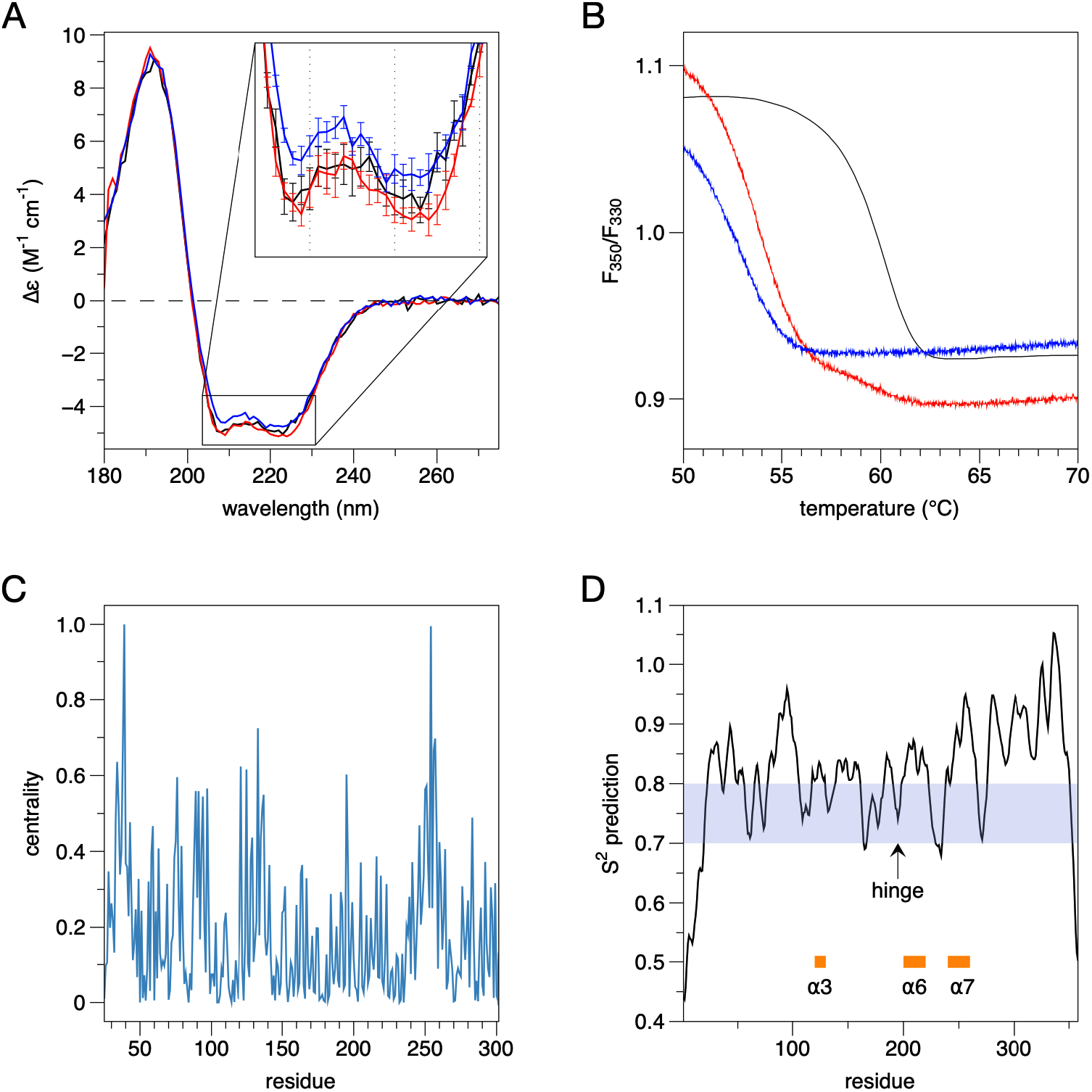
Folding and stability of GDAP1. A. SRCD spectra for wtGDAP1 (black), R120W(red), and H123R (blue). B. DSF stability assay. Colours as in A. C. Residue centrality, as defined by NAPS, identifies helix α7, around residue 250, as the most central part of the structure. D. DynaMine analysis. The location of the hinge in helix α6 is indicated, as are helixes α3, α6, and α7. The shaded region indicates context-dependent folding, while values above 0.8 predict rigid structure.

To determine thermal stability, we studied the R120W and H123R variants using the Trp fluorescence emission peak ratio at wavelengths 350/330 nm in nanoDSF. The wtGDAP1 protein is more stable than the mutants (**Fig. 3B**). The apparent T_m_ value for wtGDAP1, ~62 °C, was >5 °C higher than for both mutants, suggesting that the effect of the mutations may be linked to an overall destabilization of the fold. Considering the location of the mutations, a region of GDAP1 is revealed, which is important for protein stability.

To obtain further insight into effects of the mutations on GDAP1, we used a variety of bioinformatics tools. Analyses of centrality (**Fig. 3C**) indicated that the core region of GDAP1, close to both Arg120 and His123, with helix α7 the most central element, is likely to be important for folding and stability. Many other CMT mutations cluster into this area [18], affecting a number of residues in an interaction network (**Fig. 1A,C**). Effects of point mutations on protein stability against temperature or chemicals were predicted using CUPSAT [49]. Both R120W and H123R are predicted to be destabilizing in both respects, in line with the thermal stability data above. DynaMine analysis **(Fig. 3D**) of protein flexibility based on sequence data further showed that the part of helix α6 at before residue 200 has context-dependent rigidity, indicating that the unique α5-α6 loop in GDAP1 is structurally dynamic.

### Flexibility of the α6 helix

A crystal structure was solved for wtGDAP1Δ295-358, and it presented a novel crystal form with 4 monomers in the asymmetric unit. One complete dimer was present in the asymmetric unit, in addition to two half-dimers, which both homodimerise through crystallographic symmetry. As the resolution of the structure was rather low, structural details were not analysed. However, in all 4 independent protomers, helix α6 breaks in the middle around Asp200, and a helix at residues 189-198 is present in electron density (**Fig. 4A**). Thus, the long α6 helix can adopt different conformations even in the crystal state. The wtGDAP1 crystal structure published earlier (**Fig. 1A**) had an asymmetric dimer, with one short and one long α6 helix [18]. These observations suggest that the GDAP1-specific insertion is flexible also in solution.

**Figure 4.**
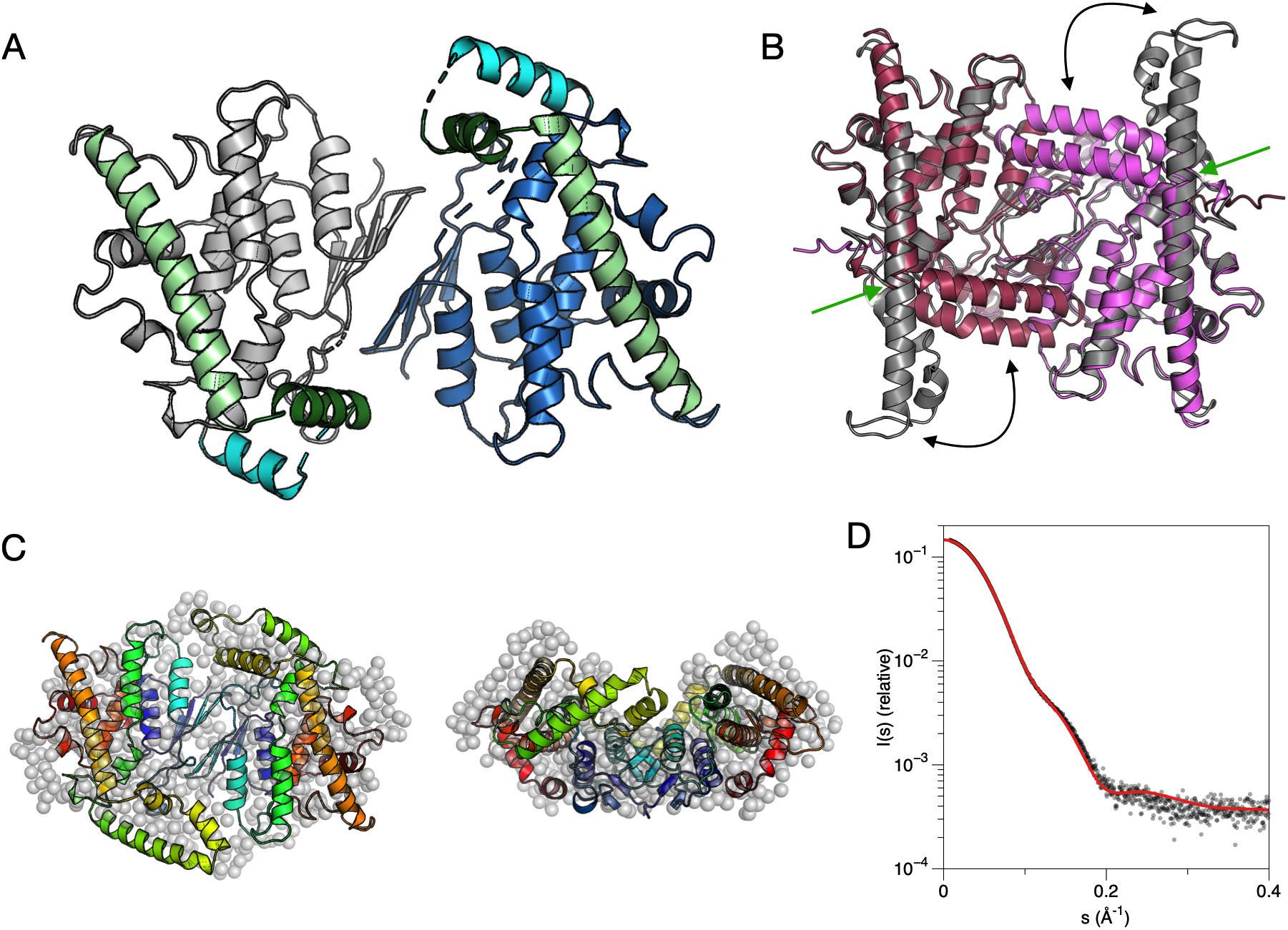
New crystal form of wtGDAP1. A. The new structure has the long α6 helix divided into two (light and dark green). B. Comparison of modelled dimers based on the extended crystal form of wtGDAP1 (gray) and AlphaFold2 (red/magenta). The double arrows indicate flexibility of the α5-α6 segment, while the green arrow identifies the hinge in the middle of α6. C. Superposition of the chain-like model with the model based on the new crystal structure (cartoons). See further comparisons in Fig. S3. D. Fit of the cartoon model from panel C to the wtGDAP1 SAXS data.

The new wtGDAP1 dimer structure was analysed with respect to the SAXS data, together with a dimer built based on the AlphaFold2 model monomer. The AlphaFold2 model has the helix α6 divided into two and collapsed into a similar, but even more compact, conformation as seen in the new wtGDAP1 crystal (**Fig. 4B**). However, it is evident that the AlphaFold2 model is too compact, while the extended conformation of helix α6 is too elongated (**Fig. 4B, S3**). An excellent fit to the SAXS data was obtained using the dimer from the new wtGDAP1 crystal structure with built-in missing loops (**Fig. 4C,D, S3**).

To further analyse the dynamics of GDAP1, the model based on the wtGDAP1 crystal structure [18], with all loops added, was subjected to MD simulations (**Fig. 5**). Throughout the simulation, the GDAP1-specific insertion is the most dynamic segment of the protein, but the long helix remains extended. On the other hand, simulation of the AlphaFold2 model indicates stability of the bent conformation. The simulations support the crystal structures of both conformations and give additional proof about a hinge in the middle of helix α6.

**Figure 5.**
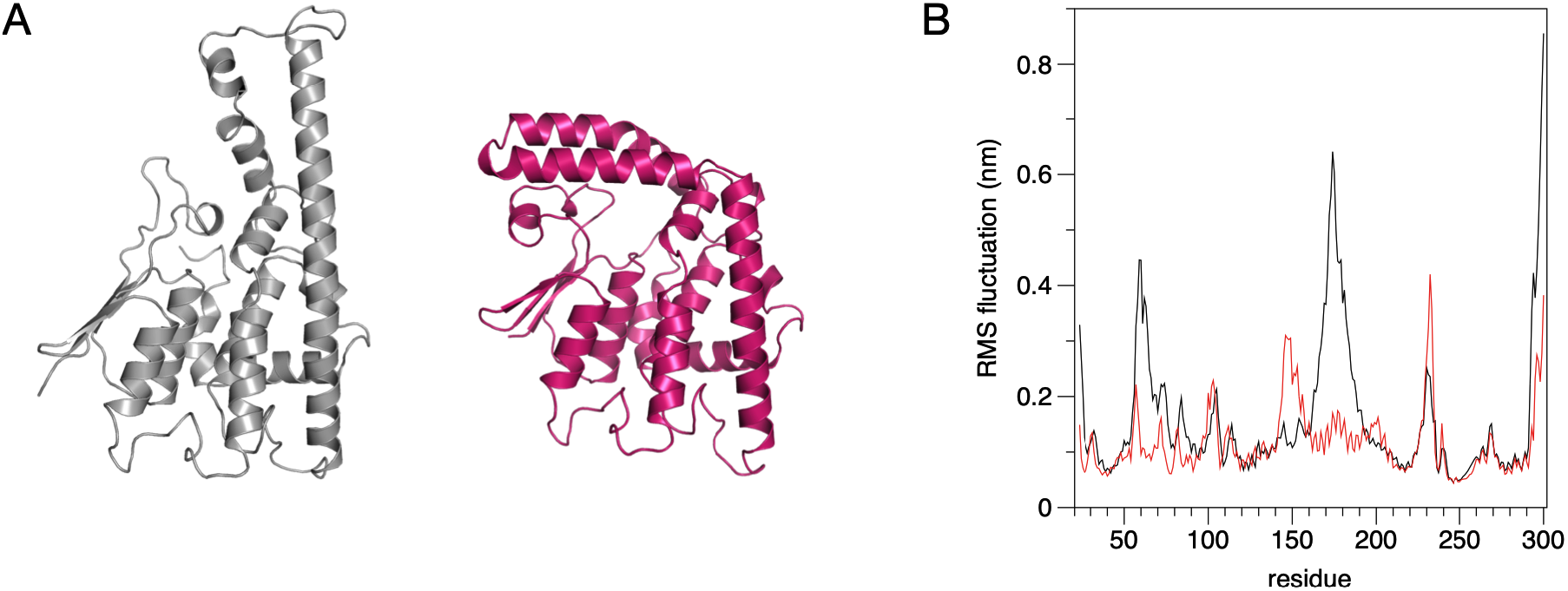
MD simulation of GDAP1 monomer. A. The open (gray) and closed (red) conformations of a GDAP1 monomer, obtained from the extended crystal structure and the AlphaFold2 model, respectively. B. RMS fluctuations of Ca atoms indicate relative rigidity of the AlphaFold2 model (red) at the β2-β3 loop and the α5-α6 loop, compared to wtGDAP1 (black). The simulations were run for 500 ns for the AlphaFold2 model and 350 ns for the open crystal structure model.

### GDAP1 mutant localization and oligomeric state in cells

To explore the two CMT-linked mutations, R120W and H123R, at the cellular level, we used three different cell culture models, in which either endogenously expressed GDAP1 variants or inducible systems to overexpress GDAP1 variants were utilized.

The localisation of R120W and H123R in neurons was compared to that of wtGDAP1 using rDRG primary cell cultures. After protein induction, the neurites were immunostained and imaged using a confocal microscope (**Fig. 6**). FLAG-tagged wtGDAP1 locates both in the cell body and axons of the sensory neurons. The localisation is not cytoplasmic or on the plasma membrane, and in accordance with previous work, most likely mitochondrial. No clear difference in the immunostaining intensity or cellular localization of R120W or H123R were observed compared to wtGDAP1. None of the mutations induced distinguishable morphological changes in sensory neurons, and no cell toxic effects were observed (**Fig. 6**).

**Figure 6.**
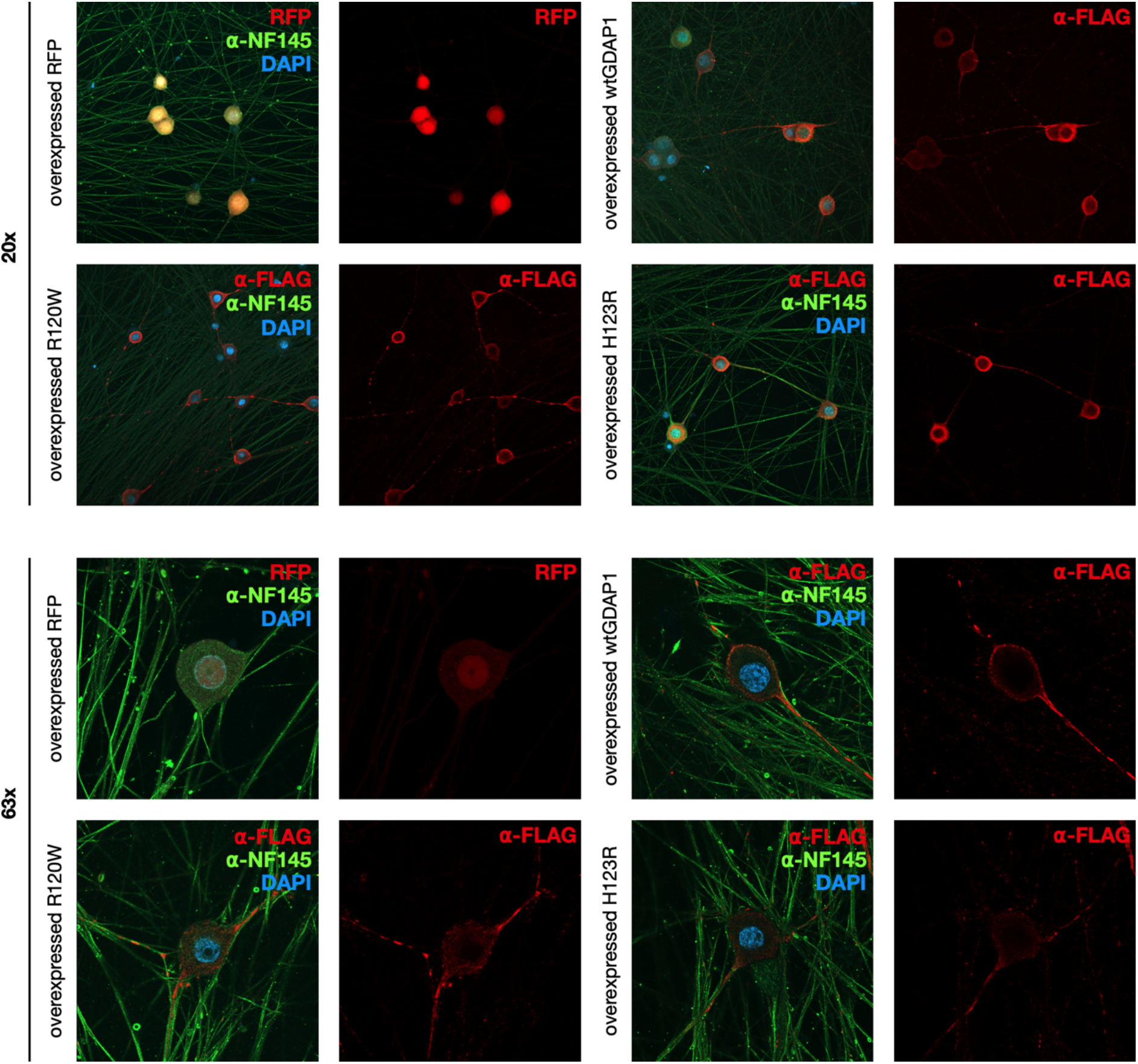
Immunofluorescence analysis of GDAP1-overexpressing rat DRGs. The images were taken with two magnifications from cells overexpressing wtGDAP1, H123R, R120W, or red fluorescent protein (RFP). GDAP1 staining was done with α-FLAG, and α-NF145 was used to visualize neurons. DAPI staining shows the nucleus.

Dimeric GDAP1 has been detectable from mammalian sources [65]. To further test whether GDAP1 preferably forms disulphide-linked dimers in cells, we performed Western blot analysis for protein extracts under non-reducing conditions **(Fig. S4).** We used human fibroblast cultures established from skin biopsies of a CMT2K patient carrying the H123R GDAP1 allele [12, 59]. Normal GDAP1 genotype fibroblasts were used as control. Western blot analysis revealed that in the fibroblasts, GDAP1 exists as monomers in both CMT2K patient-derived cells and healthy control samples (**Fig. S4).**

In addition to the human fibroblasts, we analysed the rDRG sensory neurons overexpressing GDAP1 variants and HEK293T-D10 cells that endogenously express GDAP1 to explore the oligomeric state of wtGDAP1, R120W and H123R was explored. In all cells studied here, wtGDAP1, as wells as both mutant variants, was detected as a monomer after electrophoresis (**Fig. S4**). Hence, any dimer present in the cells is not disulphide-linked. Of note, the mutation C88A did not make recombinant GDAP1 fully monomeric in our earlier study [18], indicating that the homodimer can form without the disulphide bridge *in vitro.* The only GDAP1 variant we have observed to be fully monomeric is the double mutant Y29E/C88A [18], which disturbs both the hydrophobic interface and the disulphide bridge. To conclude, the main form of GDAP1 in cells, at least in the absence of inducing factors, is not a disulphide-linked dimer.

## DISCUSSION

Single-amino-acid substitutions can alter protein physicochemical properties, affecting protein stability and function. From a clinical perspective, inherited neuropathies are generally well-characterized at the level of the symptomatic spectrum and disease progression. A large variety of CMT mutations are known and characterized clinically [66]. However, in many cases, the molecular basis of these disorders cannot be adequately explained. The difficulty of understanding the mechanism is due to both the vast number of the involved genes and their heterogeneous inheritance patterns and phenotypes, as well as limited knowledge about molecular structure and function.

*GDAP1* is one of the genes associated with peripheral neuropathies caused by missense mutations. We performed structural analyses for two human CMT2K-linked GDAP1 mutations on helix α3, which revealed that apart from the mutated residue and its immediate surroundings, the overall fold does not change. However, both mutations introduce changes in intramolecular networks and differences in molecular properties, most notably in thermal stability. The structural analysis of pathogenic CMT-linked GDAP1 variants shows that the mutations are close to the GDAP1 hydrophobic cluster and mediate interactions between key helices of the structure.

### CMT mutations in GDAP1 cluster into hotspots in 3D space

Currently, there at least 103 *GDAP1* mutations linked to CMT, out of which 68 are reported missense mutations [67]. Both R120W and H123R are common mutations in European patients [7, 14, 68], and H123R was identified as a major founder mutation in the Finnish population [12, 13]. Several GDAP1 mutations have been studied using neurons and Schwann cells or yeast models [7, 10, 69, 70]. We chose to investigate the Finnish H123R founder mutation, as well as R120W due to its well-established clinical and molecular characterization and its location in the vicinity of His123, on helix α3. Both mutations have been linked to the autosomal dominant form CMT2K.

The GDAP1 crystal structure allows predicting the molecular basis for many of the known mutations in the Human Gene Mutation Database (http://www.hgmd.cf.ac.uk/ac) and the Inherited Neuropathy Variant Browser database (https://neuropathybrowser.zuchnerlab.net). A CMT-related mutation cluster of GDAP1 (**Fig. 1**) mainly localizes on helices α3 and α6 and less on helices α7, α8, and their connecting loops [18]. There are 68 known missense GDAP1 mutations, involving 39 residues. The main cluster contains 27 residues that form a network of interactions, including salt bridges, hydrogen bonds, and van der Waals contacts. These interactions provide extensive contacts between helices α3, α6, and α7 (**Fig. 1C**). Centrality analyses of the GDAP1 structure highlight this, indicating helix α7 as the most central segment of the GDAP1 structure. Notably, many mutations linked to CMT2K map very close to each other in 3D space, suggesting an intramolecular network that gets disturbed upon disease mutations, altering GDAP1 structure or function.

Mutating residues His123 and Arg120 does not break the GDAP1 fold, but rather may affect the intramolecular residue interaction network and protein stability. The thermal stability of the mutant proteins decreased compared to wtGDAP1, suggesting that especially the interaction between α3-α6 may be important. Predictions of ΔΔG using computational methods mainly agree with the experiment, showing that both mutations are destabilising. Interestingly, the wedge between helices α5-α6 contains a pattern of double tyrosines, double glutamates, double leucines, and double lysines, which seem to pull the GSTL-C core together. The mutations in many cases are introduced into the neighbouring positions of these double pairs, like in the case of H123R.

We have shown that hexadecanoic acid bound into a groove close to the CMT mutation cluster [18]. The R120W and H123R mutation sites are close to the α6 helix and the main hydrophobic cluster centered around α7. As an interesting hotspot, Arg120 is the site for four different CMT mutations. Mutations in such clusters and hotspots might affect residue interaction networks and thus decrease protein stability.

### The cluster of interactions is sensitive towards CMT mutations

Considering the interactions of both His123 and Arg120, as well as the networks between helices α3, α6, and α7, it becomes evident that several involved residues are targets of CMT mutations. An example is Glu222, which is sandwiched between three Arg side chains (Arg120, Arg225, Arg226) and Tyr124, and has can der Waals contacts to Leu239. Another example is Ala247; the conservative mutation to valine is linked to disease [71]. Ala247 on helix α7 is located in a tight space right below His123 and Arg120 and part of the same interaction network. The apparently mild CMT mutation A247V increases the side of the side chain and affects the local interactions. Cys240 from the α6-α7 loop also lies right below His123 and Arg120, and its mutation to Tyr in CMT [72, 73] will interfere with the local interaction network at this residue cluster.

Taken together, although the CMT mutations in GDAP1 initially appear to be scattered throughout the sequence, in the 3D structure, they are involved in close interaction networks, and these networks are sensitive against changes in many different participating residues. This observation explains the general loss of protein stability upon mutations in such networks and clusters and may hint at an overall mechanism of GDAP1-linked CMT.

### The unique α6 helix of GDAP1

Helix α6 is a dominant and unique feature of the GDAP1 structure, being part of the GDAP1-specific insertion, together with its preceding loop, which is not visible in electron density. We used a combination of crystal structures and computational models to get further insights into the GDAP1 helix α6 and its dynamics. Our observation of the α6 conformational flexibility may point out to mechanistically important functions, which could be linked to effects of CMT disease mutations on or near the helix.

In the original wtGDAP1 structure [18], we saw breakdown of non-crystallographic symmetry, as the α6 helix was of different length between the two chains in the asymmetric unit. The shorter version of the helix started around residue 200, which is the hinge region in our new wtGDAP1 crystal structure, in which the helix continues in another direction in all four independent protomers in the asymmetric unit. A break of the helix at the same position is predicted also by AlphaFold2; however, the conformation of monomeric GDAP1 in the model is incompatible with the exact mode of dimerization we observe in the crystal state, leading to steric clashes. Sequence-based analysis of flexibility also points out the region around residues 190-200 as potentially flexible. The conformational dynamics of the GDAP1-specific insertion, *via* a hinge around residue 200, could play a role in its physiological functions and its interactions with other molecules, such as the cytoskeleton, *in vivo.*

### GDAP1 is a dimer in solution but not disulphide-linked in cells

GDAP1 has a unique dimer interface compared to canonical GSTs [18] and dynamic oligomerization properties [18, 65, 74]. Our results show that GDAP1 dimerization mediated *via* a disulphide bond can also be observed in the CMT mutant proteins *in vitro.* In the cellular environment, such a disulphide bond could be formed *via* a folding catalyst or through changes in the redox environment. The latter has been linked to GDAP1 function [75].

An interesting possibility for the dimeric function would be GDAP1 activation by a folding catalyst, affecting interactions with a partner protein, suggesting that the GDAP1 function could be linked to its oligomeric state. Many binding targets have been proposed for GDAP1, such as tubulin and other cytoskeletal components [23, 70]. The GDAP1 disease mutations could, in addition to affecting protein folding and stability, modulate protein-protein interactions. However, the details of molecular interactions formed by GDAP1 remain unknown, and further studies should be focused on studying GDAP1-cytoskeleton interactions and their links to GDAP1 oligomeric state.

### Conclusions

We have presented a structural analysis of two CMT-linked mutations in GDAP1, R120W and H123R. The effects of these mutations on protein structure were small, and it is likely that the mutations affect dynamic properties, stability, and conformational changes of GDAP1 and/or its interactions with additional binding partners. The cluster of CMT-related mutations in the 3D structure of GDAP1 highlights a tightly interwound network of amino acid side chain interactions that are likely essential for the normal function and structure of GDAP1. Such mutation clustering essentiates the need for accurate structural studies of proteins targeted by disease mutations, and it can be expected that most of the mutations in such a cluster similarly affect protein stability or functional interaction networks. A major goal for the future will be structure solution of the full-length GDAP1 protein, including the transmembrane domain, in addition to deciphering details of its physiological function and molecular interactions.

## Acknowledgements

This work was funded by the Academy of Finland, project number 24302881. The research leading to this result has been supported by the project CALIPSOplus under the Grant Agreement 730872, as well as by iNEXT, grant number 653706, from the EU Framework Program for Research and Innovation HORIZON 2020. We acknowledge the use of the Core Facility for Biophysics, Structural Biology and Screening (BiSS) at the University of Bergen, which has received funding from the Research Council of Norway (RCN) through the NORCRYST (grant number 245828) infrastructure consortium. The availability of synchrotron beamtime and support on DESY, EMBL/DESY, ISA, and SOLEIL are gratefully acknowledged. Furthermore, we wish to thank Dr. Roman Chrast for project planning and organization.

