## Supplementary Information for "Structural insights into Charcot-Marie-Tooth disease-linked mutations in human GDAP1"

**Supplementary Figures**

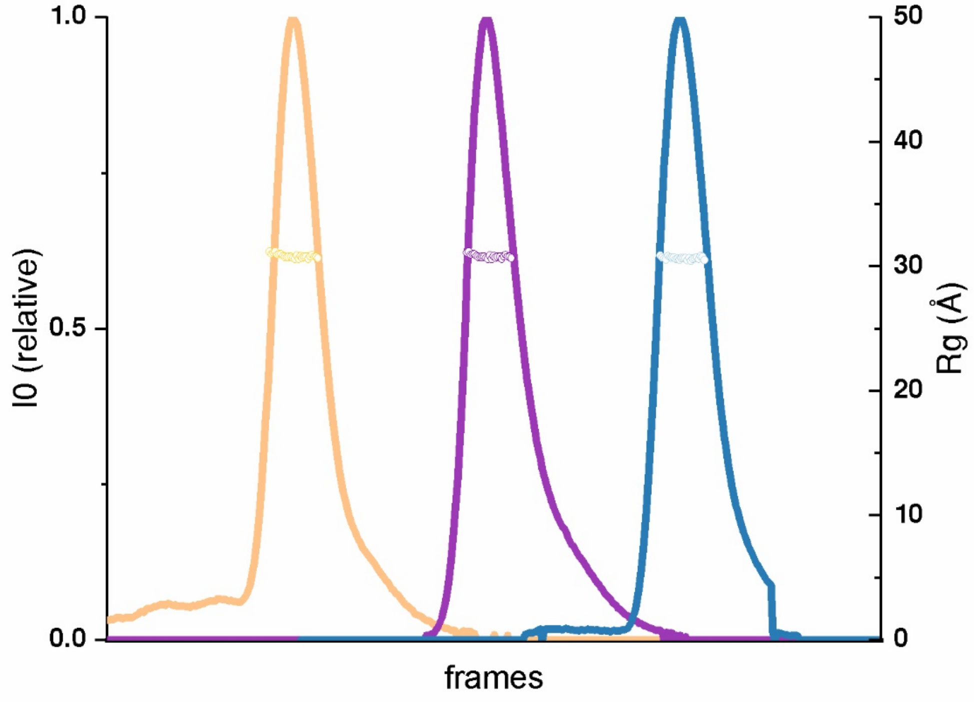


**Figure S1.** In-line SEC-SAXS frames and scattering pattern from H123R, R120W and wild-type sample analysis shown as magenta, blue and yellow respectively. Rg is shown for the parts of the peak that were processed further.

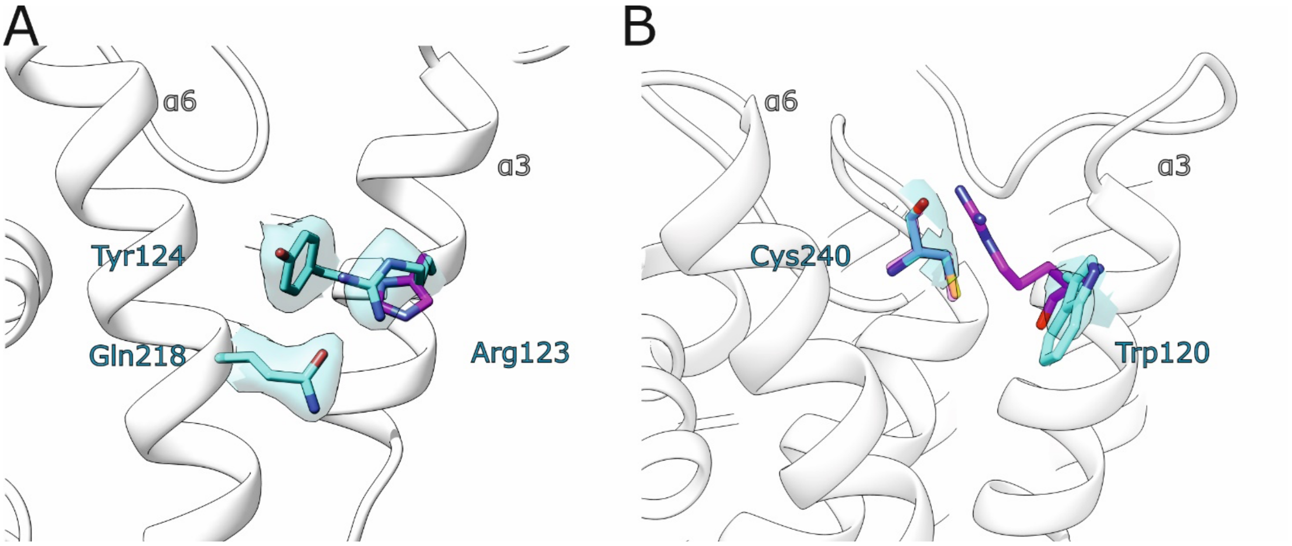


**Figure S2**. Electron density maps in the mutation sites H123R (**A**), and R120W (**B**). Both unbiased map contours were set to 1.2 rmsd, and the map radius was set to 1.6 Å from the mutated residue. Conformation of the corresponding wild-type residue is shown in magenta in each structure.

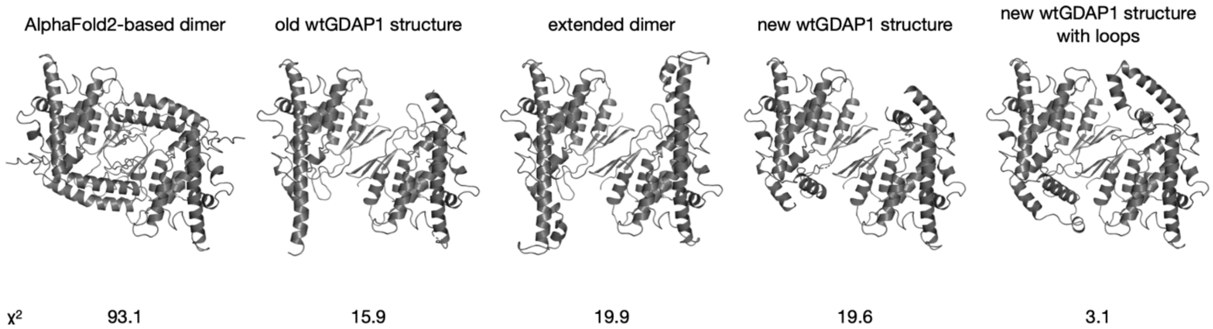


**Figure S3. Fitting of different models to the SAXS data.** Shown are models and corresponding fits to wtGDAP1 SAXS data. The best fit is obtained with the conformation from the new wtGDAP1 structure, when missing loops are built. See fit in Fig. 4D.

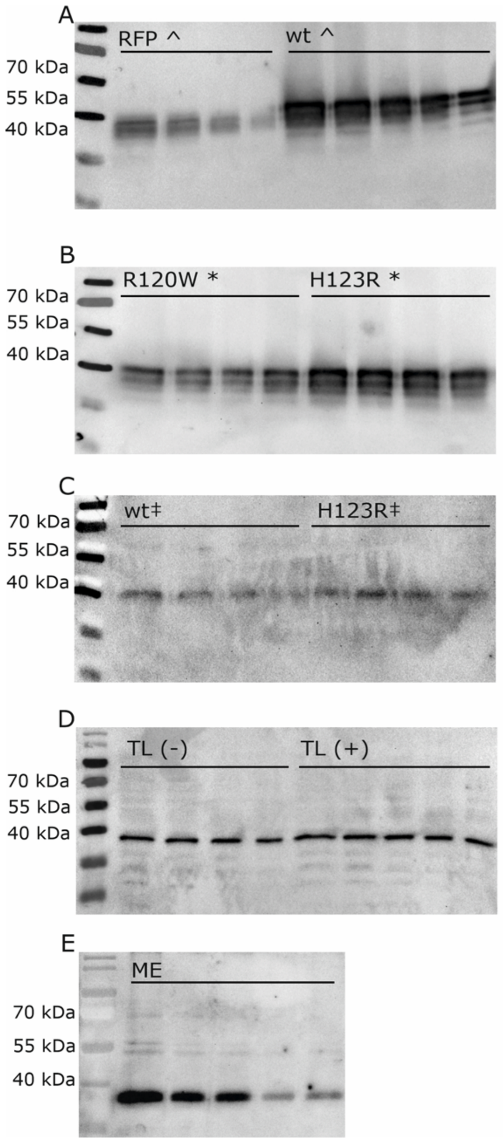


**Figure S4. Western blotting for oligomeric state of GDAP1 in cell lysates.** Western blot analysis of rat DRG sensory neurons overexpressing RFP and wtGDAP1 (A) as well as R120W and H123R (B). C. Western blot of human fibroblasts with normal (wt) and disease (H123R) genotype. D. Total lysate (TL) from HEK293T cells. The blot was run with (+), and without (-) β-mercaptoethanol in the sample buffer. E. Mitochondrial extract from HEK293T cells.
